# Machine Learning Optimization of Candidate Antibodies Yields Highly Diverse Sub-nanomolar Affinity Antibody Libraries

**DOI:** 10.1101/2022.10.07.502662

**Authors:** Lin Li, Esther Gupta, John Spaeth, Leslie Shing, Rafael Jaimes, Rajmonda Sulo Caceres, Tristan Bepler, Matthew E. Walsh

**Author notes:** These authors contributed equally to this work (Rajmonda Sulo Caceres, Tristan Bepler, Matthew E. Walsh).

## Abstract

Therapeutic antibodies are an important and rapidly growing drug modality. However, the design and discovery of early-stage antibody therapeutics remain a time and cost-intensive endeavor. In this work, we present an end-to-end Bayesian, language model-based method for designing large and diverse libraries of high-affinity single-chain variable fragments (scFvs). We integrate target-specific binding affinities with information from millions of natural protein sequences in a probabilistic machine learning framework to design thousands of scFvs that are then empirically measured. In a head-to-head comparison with a directed evolution approach, we show that the best scFv generated from our method represents a 28.8-fold improvement in binding over the best scFv from the directed evolution. Additionally, 99% of the designed scFvs in our most successful library are improvements over the initial candidate scFv. By comparing a library’s predicted success to actual measurements, we demonstrate our method’s ability to explore tradeoffs between library success and diversity during the design phase and prior to experimental testing. The results of our work highlight the significant impact machine learning models can have on scFv development. We expect our end-to-end method to be broadly applicable and able to provide value to other protein engineering tasks.

## Introduction

Therapeutic antibodies are an important and rapidly growing drug modality. Because the vast search space of antibody sequences renders exhaustive evaluation of the entire antibody space infeasible, screening relatively small numbers of antibodies from synthetic generation, animal immunizations or human donors is used to identify candidate antibodies. The screened library represents a small portion of the overall search space, and the resultant candidate antibodies are often weak binders or suffer from developability issues. Optimization of these candidates is needed to improve binding and other development characteristics.

Due to the combinatorial scaling of sequence space, step-wise, iterative approaches are often used to optimize antibody binding against target molecules [1], [2], but are time consuming and effort is wasted interrogating nonfunctional antibodies. Improved binders may need to be further altered to improve other properties, such as hydrophobicity [3], [4], but such alterations can negatively influence the previously optimized binding, resulting in additional measurement and engineering cycles. This process of identifying the final antibody routinely takes about 12-months to complete [2]. The ability to efficiently engineer antibodies with favorable binding and high diversity earlier in the development process would reduce the impact of unfavorable antibody characteristics that are often identified later in the process, improve the developability potential and reduce the time required in early drug development.

While computational methods can guide the search of biologically relevant antibodies, most *de novo* approaches require target structures or antibody-epitope complex structures to be known [5]–[7]. Machine learning (ML) approaches can be used to effectively represent biological data and rapidly explore their vast design spaces in silico. Such approaches can uncover complex and flexible features from high-dimensional data [8]–[13] and have shown great promise in many application areas, including protein structure prediction [14], and drug discovery and design [15]–[20]. Similarly, existing ML-driven antibody optimization has shown promising results in designing antibodies with improved binding characteristics against a target and that antibody binding can be learned from only sequence data and without the need for the target’s structure [15]. A more recent work has presented an ML-driven antibody optimization approach that achieves broader neutralizing activity against diverse SARS-CoV-2 variants by learning the mutational effect on protein-protein interactions from protein complex structures [20]. Other works have investigated general purpose pre-trained generative language models for designing antibody libraries that display good physical properties [18], [19], but these methods are not target-specific and only offer modest improvements over conventional libraries that are, often, already based on natural antibody repertoires. Finally, none of the existing work allows the evaluation of designed antibody libraries prior to experimentation, a critical feature that allows for accelerated design cycles.

In this work, we develop an end-to-end ML-driven single-chain variable fragment (scFv) design framework that uniquely combines state-of-art language models, Bayesian optimization and high-throughput experimentation (Fig. 1). Because we synthesize explicitly defined oligo pools of 200bp, our method allows the design of the entire scFv chain (heavy or light). Furthermore, it does not assume candidate scFvs strongly bind to the target, and relies on sequence data without the need for sequence alignments or knowledge of the target antigen structure, allowing the method to be applicable to early-stage antibody development for any target antigen. We demonstrate our end-to-end framework can rapidly and cost-effectively lead to the design of diverse target-specific scFv libraries with therapeutically relevant binding affinities. At a meaningful scale (~10^4^ sequences), and in a head-to-head comparison with the directed evolution approach, we show that our ML-based approach produces significantly stronger binders. More remarkably, our ML-designed scFv libraries are highly diverse, demonstrating the ability of our approach to efficiently extrapolate and discover mutationally distant, high affinity scFvs. Lastly, we show how our method can provide general insights to the engineering process. We can evaluate the performance of an scFv library in silico, explore the affinity-diversity tradeoff prior to experimental testing, weigh the choice of optimizing CDRs jointly or individually, and combine our method with other software tools to explore other desired development properties, such as hydrophobicity and isoelectric point, of scFvs in designed libraries. Our results highlight the impact ML models can have on early-stage scFv development. Through coordinated data generation, ML model development, training and optimization, we are able to start with only a target protein sequence and after a single round of optimization, generate large, diverse libraries of high-affinity scFvs against the target.

**Fig. 1.**
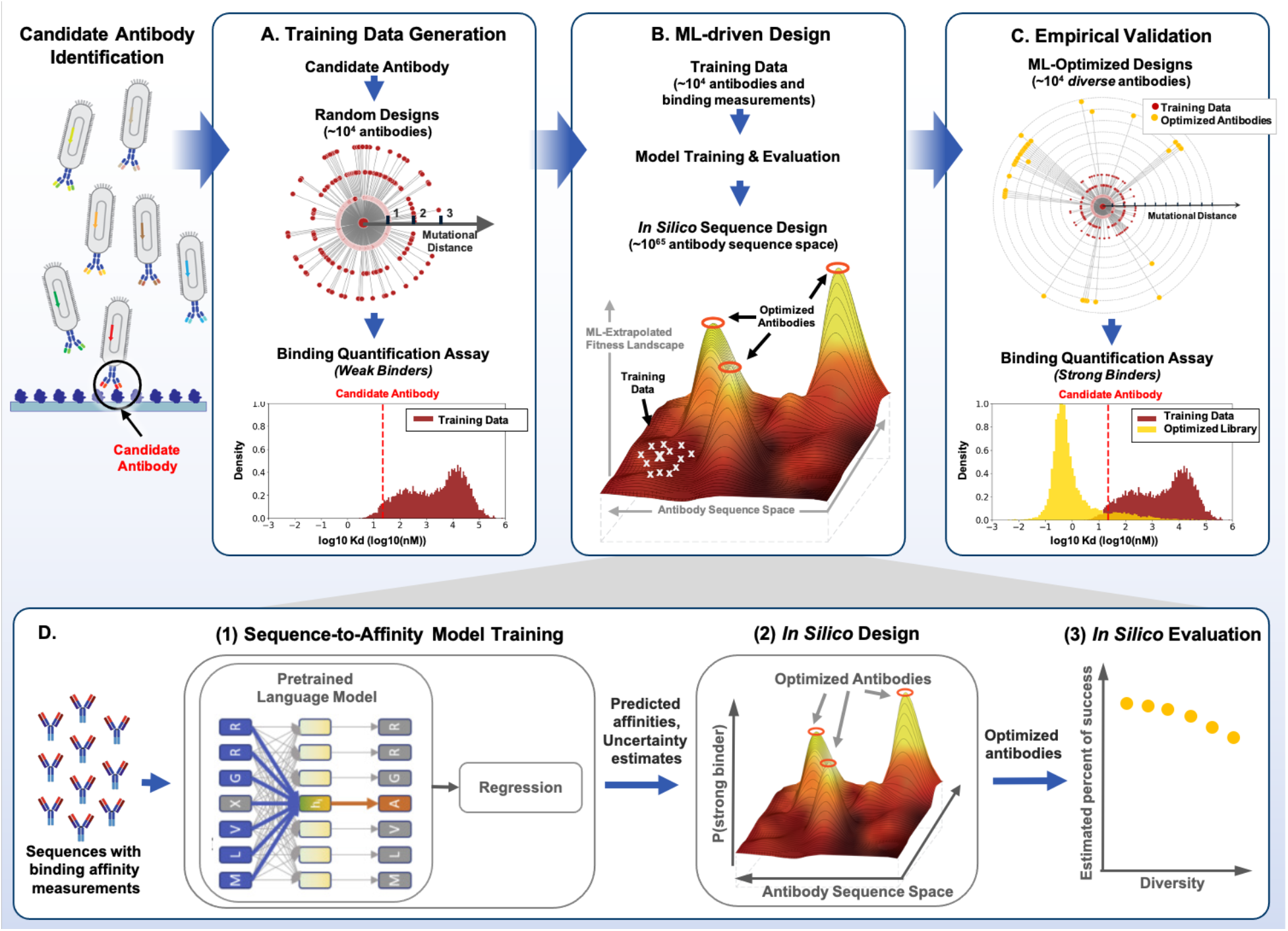
Illustration of the end-to-end ML-driven antibody design process. **A**. The training data is generated via random mutations of the candidate scFv antibody along the entire CDR region, followed by high-throughput binding quantification. **B**. This training data combined with publicly available protein sequences is used to train, refine and evaluate ML models that drive the in silico sequence design process. **C**. The designed libraries are experimentally validated. ML-driven designs produce highly diverse antibodies (sequences as far as 23 mutations away), with strong on-target binding (the best design is 2818% better than the directed evolution approach), and high success rate (as high as 99%). **D**. Detailed ML-driven design process: (1) supervised fine-tuning of pretrained language models on the training data to predict binding affinities with uncertainty quantification; (2) in silico scFv antibody design via Bayesian optimization over ML-extrapolated fitness landscape; (3) in silico antibody library evaluation

## Results

### Development of an end-to-end, target-specific scFv optimization process

We hypothesized that by integrating target-specific binding affinities with information from millions of natural protein sequences in a probabilistic machine learning framework, we could rapidly engineer scFvs that are significantly stronger binders than what typical directed evolution approaches would produce. To engineer a given candidate scFv against the target molecule, we developed a five-step process that uniquely combines state-of-art language models, Bayesian optimization and high-throughput experimentation to generate high-affinity scFv libraries (Fig. 1 and Methods section):

1. High-throughput binding quantification of random mutants of the candidate scFv to create supervised training data (Fig. 1A).
2. Unsupervised pre-training of language models [21], [22] on large numbers of protein sequences to distill biologically relevant information and represent scFv sequences (Fig. 1B and 1D).
3. Supervised fine-tuning of pretrained language models on the training data to predict binding affinities with uncertainty quantification (Fig. 1B and 1D).
4. Construction of Bayesian-based scFv fitness landscape extrapolated from the trained sequence-to-affinity model, followed by in silico scFv design via Bayesian optimization and in silico design validation (Fig. 1B and 1D).
5. Experimental validation of top scFv sequences that are in silico predicted to have strong binding affinities (Fig. 1C).

We generated our supervised training data using an engineered yeast mating assay and have published it separately to support its reuse [23]. This data includes 17,118 heavy chain scFv binding measurements and 26,223 light chain scFv binding measurements against a single target peptide and were collected in pooled, yeast-based mating assays. All heavy- and light-chain sequences in this data were generated by performing random k=1,2,3 mutations of the candidate antibody Ab-14 along the entire CDR within its respective chain. The binding measurements are provided on a log-scale, with lower values indicating stronger binding.

We pre-trained four BERT masked language models, i.e., a protein language model, an antibody heavy chain model, an antibody light chain model and a paired heavy-light chain model. The protein language model was trained on the Pfam data [33], and antibody-specific language models were trained on human naïve antibodies from the Observed Antibody Space (OAS) database [34].

To train sequence-to-affinity models, we investigated two approaches to predict affinities with uncertainty quantification: an ensemble method and Gaussian Process (GP). Both approaches use learned knowledge from pre-trained language models and provide meaningful sequence-to-affinity models from which one can design high-affinity scFv libraries. We trained separate sequence-to-affinity models for Ab-14-H heavy-chain variants and Ab-14-L light-chain variants using the corresponding training data. We observed strong positive correlation between predicted and experimentally measured binding affinities on the hold-out test data (Fig. 4A).

To generate high-affinity scFv libraries, a Bayesian-based fitness landscape was constructed to map the entire scFv sequence to a posterior probability, i.e., the probability that the estimated binding affinity is better than the candidate scFv Ab-14. This is in contrast to the fitness landscape that goes directly from sequence to estimated binding affinity. To perform optimization to maximize the posterior probability, the choice of sampling algorithm is critical in determining the library diversity. Three strategies were used: hill climb (HC), genetic algorithm (GA) and Gibbs sampling. HC is a greedy algorithm that performs a local search and only finds local maximums. GA is an evolutionary-based algorithm that is more robust in exploiting the sequence space further away from the initial sequence. Gibbs sampling takes sequential actions in a manner that balances exploitation and exploration and can generate sequences with high diversity.

We applied our sampling approaches to generate heavy chain and light chain variant scFvs that optimize Ab-14. We also used a Position-Specific Score Matrix (PSSM)-based method representative of traditional directed evolution approaches to generate a control sequence set. The generated sequences from each method are rank-ordered based on the posterior probability and top sequences are selected. This resulted in seven scFv libraries per chain: three libraries from optimizing the ensemble-based fitness function (namely, En-HC, En-GA and En-Gibbs), three libraries from optimizing the GP-based fitness function (namely, GP-HC, GP-GA, GP-Gibbs), and one PSSM library. As a sanity check, we also generated scFv mutants with an average of k=2 random mutations from the 10 strongest binders of the supervised training data. All sequences were synthesized and experimentally tested using the same high-throughput yeast display method as for the training data generation; Supplementary Table 5 and 6 provide the exact number of sequences from each library.

### ML-generated ScFv Libraries Outperform Conventional Directed Evolution

We assessed the quality of each ML-derived scFv library by comparing the binding strength of the best design and the percent of success to the PSSM-generated library. We define the percent of success as the percent of scFvs that have a better binding score than the initial scFv, Ab-14. We chose PSSM libraries as comparators because they better reflect the traditional optimization process and are generally better than random mutation libraries (Supplementary Fig. 5). **Table 1** contains characterization of the best binding scFv from each library. The sequences of these scFvs can be found in the supplementary material (Tables 7 and 8). The best scFvs from ML-optimized libraries are significantly stronger binders than those from the PSSM library, and generally have more mutations. The strongest binding heavy chain design is from the En-Gen library and binds 28.8-fold stronger than the strongest scFv in the PSSM library. The best light-chain design is in the En-Gibbs library achieving a 7.8-fold improvement over the best scFv from the PSSM library. Note that the best heavy-chain scFv binds a lot stronger to the target than the best light-chain scFv. To investigate further, we rank-ordered all designed scFvs across different libraries by the empirically-measured binding affinity and observed that heavy-chain designs are generally stronger binders than light-chain designs (Supplementary Fig. 10A).

**Table 1.** Characterization of the top scFv from each library.

| Library | Best Ab-14-H Variant Design |  |  | Best Ab-14-L Variant Design |  |  |
| --- | --- | --- | --- | --- | --- | --- |
|  | Predicted Affinity (pM) | Mutational Distance to Ab-14 | Fold Improvement Over PSSM | Predicted Affinity (pM) | Mutational Distance to Ab-14 | Fold Improvement Over PSSM |
| PSSM | 109.65 | 4 | 1.0 | 112.20 | 3 | 1.0 |
| GP-HC | 52.48 | 3 | 2.1 | 57.54 | 3 | 1.9 |
| GP-GA | 20.42 | 4 | 5.4 | 16.60 | 3 | 6.8 |
| GP-Gibbs | 15.49 | 4 | 7.1 | 100.00 | 9 | 1.1 |
| En-HC | 3.80 | 7 | **28.8** | 154.88 | 11 | 0.7 |
| En-GA | 3.89 | 10 | 28.2 | 30.20 | 17 | 3.7 |
| En-Gibbs | 38.01 | 15 | 2.9 | 14.45 | 23 | **7.8** |

Fig. 2 shows the performance and diversity of designed libraries. For Ab-14-H heavy chain designs, with the exception of sequences in the En-Gibbs library, all ML-optimized libraries outperform the PSSM library in terms of median binding affinity (Fig. 2A), and are significantly more successful than the 23.8% success of the PSSM library (Fig 2B). The En-HC (94.3%) and En-GA (96%) libraries are particularly successful and outperform all GP-generated libraries (59.4% - 84.2%). For the Ab-14-L light chain designs, all ML-optimized libraries outperform the PSSM library in both median binding (Fig. 2D) and percent of success where the PSSM library is 45.6% successful (Fig. 2E). The percent of success of GP-based libraries (95.7-99%) further outperforms all ensemble-based libraries (67.9-73.5%).

**Fig. 2.**
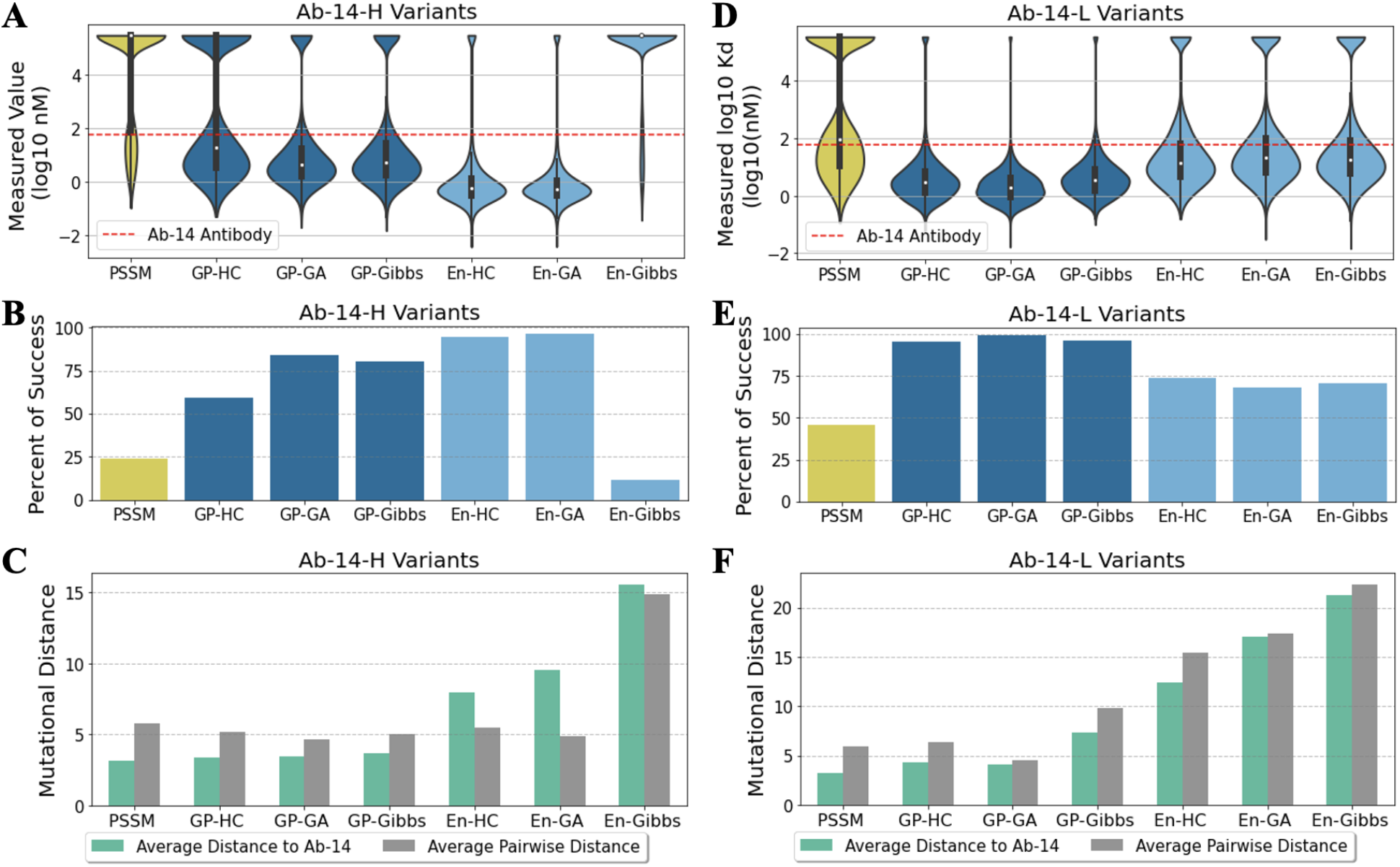
ML-optimized antibody libraries outperform the PSSM directed evolution approach with high percentage of success and high diversity. The binding quantification assay was conducted from six technical replicates. A replicate with an empty measured affinity value indicates that it was beyond the limit of detection and was deemed poor binders. For sequences that have at least 3 (out of 6) empirical binding affinities, we use the averaged values as ground-truth measured affinities. The rest of the sequences (with less than 3 empirical measurements) are automatically considered as un-successful designs. (A) Measured affinity distribution of Ab-14-H heavy chain designs (excluding sequences with no ground-truth affinity value). Affinities of unsuccessful sequences are set to be 5.48 (the largest assay value of all Ab-14-H variants). (B) Percent of sequences that have stronger binding affinity than the candidate antibody for all the Ab-14-H variant libraries. (C) Diversity comparison for all the Ab-14-H variant libraries. (D) Measured affinity distribution of Ab-14-L light chain designs. Affinities of unsuccessful sequences are set to be 5.53 (the largest assay value of all Ab-14-L variants). (E) Percent of success for all the Ab-14-L variant libraries. (F) Diversity comparison for all the Ab-14-L variant libraries.

### ML-generated Libraries can be Highly Diverse

We measured the library diversity using two mutational distance metrics: 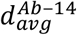 (the average distance to the initial Ab-14), and *d_pw_* (the average pairwise distance). The former 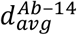 indicates how far the designs are from the training data and the latter *d_pw_* measures the intra-library diversity. For Ab-14-H variant designs, all ML-optimized libraries have higher 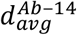 than the PSSM library (with 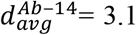). The ensemble-based libraries also have significantly higher 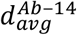 (7.9 - 15.6) than the GP-based libraries (3.4-3.7), indicating that the methods are able to extrapolate and design sequences that are far beyond the training data (Fig. 2C). In particular, sequences in the En-Gibbs library are on average 15.6 distance away from Ab-14-H and 14.9 distance away from each other (Fig. 2C). However, this increase in mutational distance comes at the cost of reduced affinity, suggesting that there is eventually a tradeoff between the two.

For Ab-14-L variant designs, all ML-optimized libraries are significantly further away from Ab-14-L than the PSSM library, with 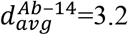 for the PSSM library, 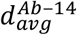 ranging from 4.3 to 7.4 for GP-based libraries and 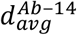 ranging from 12.4 to 21.3 for ensemble-based libraries (Fig. 2F). With the exception of GP-GA (*d_pw_*=4.5), all ML-optimized libraries have higher *d_pw_* (ranging from 6.3 to 22.4) than the PSSM library (*d_pw_*=5.9). In particular, the En-Gibbs library consists of sequences that are on average 21.3 distance away from Ab-14-L and 22.4 distance away from each other (Fig. 2F).

Fig. 3 shows the 2-D embeddings of all antibody libraries and the training data. We observed a similar trend for both light- and heavy-chain designs, that is, the PSSM library is the closest to the training data while the ensemble-based libraries are the farthest away from the training data. More interestingly, all optimization-based libraries occupy a distinct subspace from the training data and PSSM library, highlighting the extrapolating power of the various optimization approaches that we applied. Ensemble-based libraries are highly divergent and also group distinctly from the other libraries; both the best heavy- and light-chain designs were discovered via optimizing the ensemble-extrapolated fitness function, underlining the value of exploring further away from the initial candidate sequence.

**Fig. 3.**
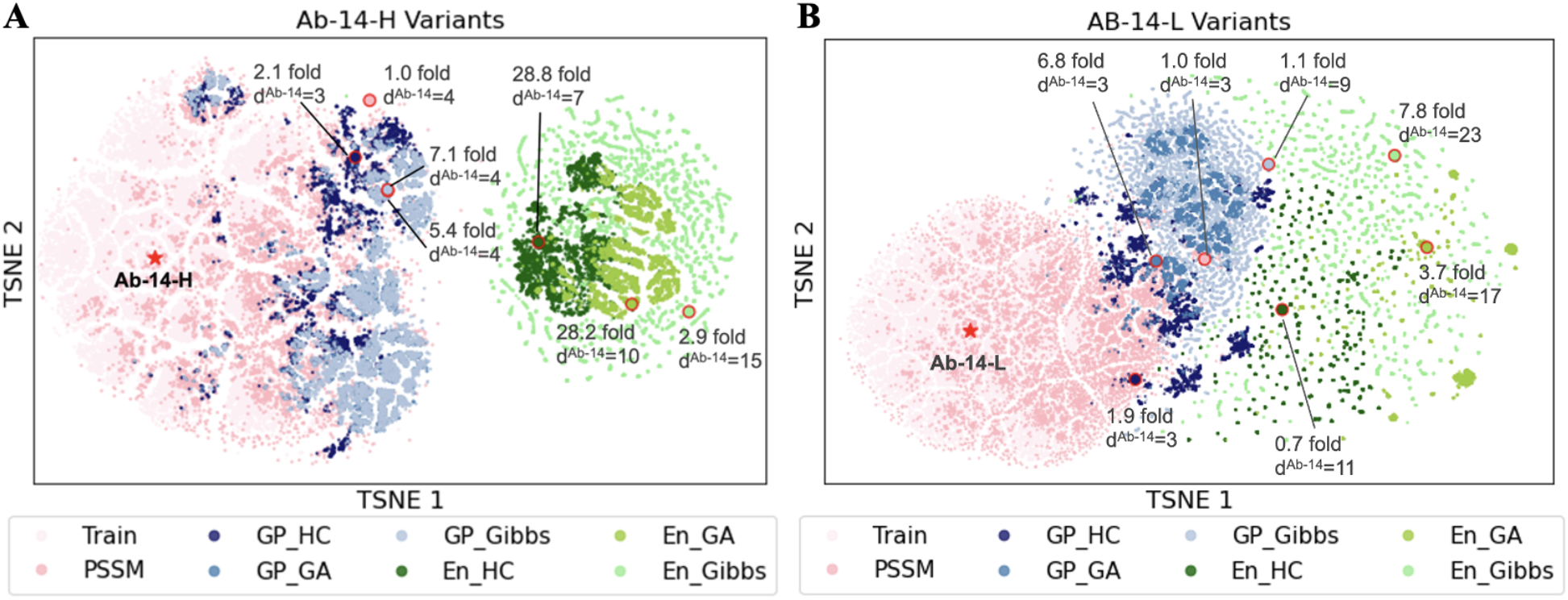
T-SNE sequence embedding of ML-libraries, PSSM library and the training data reveals distinct sampled sequence subspaces. The t-SNE embeddings allow visualization of the sequence space by embedding the sequences into 2-D based while approximately preserving the edit distance between sequences. (A) 2-D embeddings of Ab-14-H variant designs and best scFvs from each library along with fold improvement over the best scFv of the PSSM library and mutational distance from Ab-14. (B) 2-D embeddings of Ab-14-L variant designs and best scFvs from each library.

### Model Performance and Sampling Diversity are Key Factors in Generating a Quality Library

To understand key factors that determine the quality of a generated library, we first evaluated the performance of the two sequence-to-affinity models, using held-out test data and empirical binding measurements of our designed sequences (Fig. 4). We compared the Spearman correlation and the mean absolute error (MAE) of model predictions and measured values. We observed that the ensemble sequence-to-affinity model does better at predicting affinity than the GP model. When evaluated on the held-out test data, Spearman correlation scores of both heavy- and light-chain ensemble models are slightly higher (heavy-chain model: 0.51; light-chain model: 0.69) than the respective GP models; see Fig. 4A. When evaluated on designed Ab-14-L variants, the light-chain ensemble model is also slightly better with a Spearman correlation of 0.61. The most notable difference is when evaluating on designed Ab-14-H variants, where the heavy-chain ensemble model has a Spearman correlation of 0.70 but the heavy-chain GP model performs significantly worse (−0.45). This is primarily due to the prediction limit of the GP model on sequences that are far beyond the training data. When evaluated the MAE of our prediction models with respect to the mutational distance on designed sequences, we observed a sharp increase in MAE on sequences with six or more mutations away from Ab-14-H for the heavy-chain GP model, and on sequences with ten or more mutations away from Ab-14-L for the light-chain GP model (Fig 4B). Ensemble models exhibit no notable increase in MAE as the mutational distance increases, indicating that the ensemble approach is more generalizable to higher-order mutants than the GP model. Nevertheless, GP-based libraries, when compared to the PSSM library, are significantly more successful while having comparable sequence diversity (Fig. 2).

**Fig. 4.**
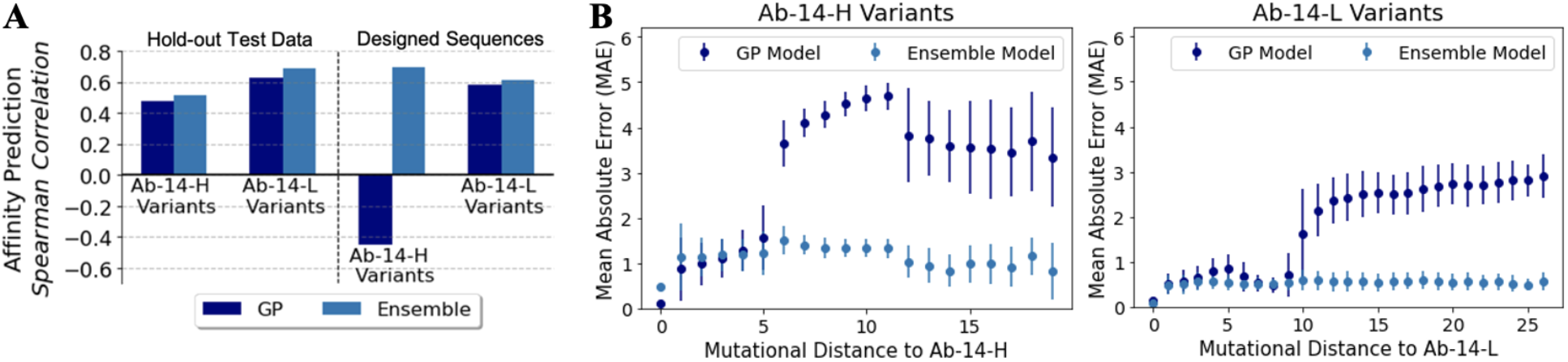
Sequence-to-affinity model evaluation. (A) Regression performance on hold-out test data and on the designed libraries; the ensemble model is more predictive than the GP model on both datasets. (B) Mean absolute error (MAE) and variance of the GP and ensemble models with respect to mutational distance from Ab-14. Evaluation was performed on sequences with at least 3 (out of 6) empirical binding affinities and the averaged values are used as the ground-truth. Ensemble models are more robust at extrapolating mutationally distant scFvs while the GP models do not predict well on sequences that are mutationally far away from Ab-14. Note that the error bar of the heavy-chain ensemble model shows a non-trivial increase on sequences that are twelve or more mutations away from Ab-14, suggesting that the model’s predictability decreases with increase in mutational distance.

While ML-guided exploration of sequence space allows for identification of more scFvs with optimized binding, we postulate that if this set comes from diverse sequence space, it will also have diverse development properties thus limiting the chance of correlated downstream development failure. We observed that a good prediction model is necessary but not sufficient to generate a diverse library with high affinity. Equally important to the prediction model is the choice of sampling algorithm. When using the ensemble-extrapolated fitness landscape to engineer 14-Ab-H, hill climb and genetic algorithms found scFvs with significant (28.8 and 28.2-fold, respectively) increases in binding over the best PSSM-sampled scFv (**Table 1**), and both methods were highly successful (94.3% and 96% success, respectively); see Fig. 2A and 2B. However, when combined with the Gibbs sampling algorithm, the best scFv sampled was only 2.9-fold better (**Table 1**), and the library was generally unsuccessful (Fig 2A and 2B). With the two diversity metrics of the En-Gibbs-generated sequences almost double that of the En-HC and En-GA libraries, it indicates that the significant increase in diversity of the En-Gibbs library has a detrimental effect on library affinity due to the eventual limit of the model predictability on sequences that are deemed too far from the training data (Fig. 2C, Supplementary Fig. 2). Interestingly, when engineering the light chain (14-Ab-L), the En-Gibbs combination found the strongest binder (7.8-fold improvement over PSSM) with a striking 23 mutations from the Ab-14-L sequence **(Table 1)**. For the ensemble-based libraries, as the library diversity increased, so too did the binding strength of its top scFv (Fig. 2F and **Table 1)**. En-HC, the least diverse ensemble-generated 14-Ab-L library, was the only library that failed to contain an scFv outperforming the top PSSM-generated scFv **(Table 1)**. In this instance, the increased library diversity is beneficial, suggesting the value in exploring away from the initial candidate sequence. To avoid unsuccessful library designs while still being able to explore sufficiently high orders of mutants, it is important to control the diversity of sampled sequences via parameter tuning of the sampling algorithm and have the ability to explore the tradeoff between performance and diversity in silico prior to experimental testing.

### Bayesian-based Approach Provides Insights Prior to Experimental Testing

We defined an in silico performance metric that quantifies the binding performance of a library prior to experimental testing. With the Bayesian approach, the fitness score is the posterior probability of a sequence in the library having a stronger binding affinity than the candidate scFv Ab-14. We average the individual fitness scores of the full library to come up with our metric - an estimate of the probability of success (i.e., the estimated percent of sequences having a better binding performance than the threshold value). We first evaluated the utility of the metric on the hold-out test data from the training scFv library as we vary the threshold value that defines strong binders and show the estimated percent of success matches well to the actual percent of success (Supplementary Fig. 7).

We applied the metric (estimated percent of success) to the designed libraries and ranked them. We compared the library ranking based on the estimated and measured percent of success (Supplementary Table 9). For PSSM and ensemble-based libraries, the predicted rankings match well to the actual rankings with a rank correlation of 0.8. For ranking PSSM and GP-based libraries, the metric predicts all rankings correctly for Ab-14-H variant libraries and a rank correlation of 0.8 for Ab-14-L variant libraries. Moreover, we observed that the estimated percent of success captures well the relative performance of designed libraries for both heavy- and light-chain designs (Supplementary Fig. 8 and 9).

We then sought to extend the application of the in silico metric to comparing the choice of optimizing one CDR to optimizing all three simultaneously. For this comparison, designs were generated using the genetic algorithm sampling over the ensemble-extrapolated fitness landscape. We observed that designing all heavy-chain CDRs leads to sequences with higher estimated percent of success than when designing individual CDRs (Supplementary Fig. 10B).

Based on these findings, we demonstrate that the performance metric can be used to understand design choices and explore tradeoffs between performance and diversity to inform library selection prior to experimental testing.

## Discussion

We demonstrate for the first time, in a head-to-head comparison with a conventional directed evolution strategy, scFvs designed with our ML approach are significantly stronger binders, especially at high levels of diversity, where, remarkably, our models are able to accurately predict binding affinity for extremely high order mutants. The libraries generated through our method have diverse biophysical properties (Supplementary Fig. 11). This allows for the selection of multiple preclinical candidates, uncorrelated in their downstream failure modes, such that if one fails, the entire pipeline is not likely to fail for the same reason. We also believe that our framework is applicable to any task aiming to maximize or minimize a characteristic of an scFv, such as minimizing off-target binding or maximizing neutralization. Pending data availability, we see ML-based multi-objective scFv optimization as an approachable task and viable option for streamlining scFv development.

We separately explored our model performance as a function of the amount of training data and demonstrated additional data, expectedly, results in improved performance [26]. However, after about 7,000 measurements, additional measurements result in less significant performance increases [26]. For this work, we trained our supervised sequence-to-affinity models on all 43,341 measurements that were available to us, but future engineering attempts may optimize use of financial resources by increasing the number of cycles while reducing the number of measurements per cycle. Because of cost limitations associated with DNA synthesis, we chose to generate our training data by introducing k = 1, 2, or 3 random mutations, but our models are able to successfully extrapolate much further than that. Future work would benefit from an improved understanding of the way in which training data is generated, if there is dependence on the choice of model, and if performing multiple measurement cycles impacts the choice.

Recently, other works have presented general purpose pre-trained generative language models for antibody design [18], [19]. By training on natural antibody repertoires, Shin *et. al*. [22] were able to design antibody libraries that display good physical properties and are enriched for binders. In the future, our approach can be combined with these by using a pre-trained generative language model to design the initial mutagenesis library used for training our supervised learning approach. Our initial analysis indicates that this approach is likely to increase the success rate of the initial library by several fold. Furthermore, pre-trained models could also condition on features of the target epitope to design target-specific initial libraries that are then fine-tuned with our framework.

We demonstrate the ability to rapidly design large libraries of potently binding scFvs, but our framework also extends to other domains of protein engineering where large scale functional mutagenesis screens are being applied. Our framework is neither scFv nor binding-specific and, therefore, can be applied to engineer other proteins for other functional properties. We expect machine learning approaches like ours, combined with high throughput mutagenesis screens, will soon become the standard in protein engineering.

## Online Methods

### Training data for Language Models

We used sequences from Pfam and Observed Antibody Space (OAS) databases to train four separate language models (i.e., a protein language model, an antibody heavy chain model, an antibody light chain model and a paired heavy-light chain model). The Pfam is a database of curated protein families containing raw sequences of amino acids for individual protein domains [24]. We use the same data splits as provided in [8]. The train, validation and test splits contain 32,593,668, 1,715,454 and 44,311 sequences, respectively. The full OAS database contains immune repertoires from over 75 studies containing a diverse set of immune states [25]. We curated only studies with naïve human subjects and removed redundant sequences across the studies. This results in 37 studies containing 270,171,931 heavy chain sequences, 9 studies containing 70,838,791 light chain sequences, and 3 studies containing 33,881 heavy-light sequence pairs. The train, validation and test sequences are split based on studies. Given that there are limited heavy-light sequence pairs in the OAS data, to train the paired heavy-light chain model, we used all the data from OAS heavy chains, OAS light chains and OAS heavy-light sequence pairs. Supplementary Table 4 summarizes the number of sequences in train, validation and test data for the four language model training datasets.

### Training BERT Language Models

We used the BERT masked language model to encode protein/antibody sequences. It models the probability of an amino acid sequence *p*(*x*) by considering the probability distribution over each amino acid at each position conditioned on all other amino acids in the sequence, that is, 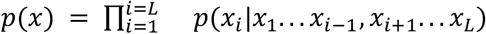, where *x_i_* represents the *i^th^* amino acid in the sequence of length L. We pretrained four separate BERT language models, i.e., a protein language model, an antibody heavy chain model, an antibody light chain model and a paired heavy-light chain model, using the Pfam data [24] and OAS data [25]. Specifically, BERT masked language models were trained with 768 input embedding size, 24 hidden layers, 1024 hidden size, 4096 intermediate feed-forward size and 16 attention heads. All the other architecture details are fixed to their default values as provided in [8]. We trained the language model to predict randomly masked amino acids in a single sequence or a sequence pair. For training the protein language model, antibody heavy chain model and antibody light chain model, the input is a single sequence of amino acids. For training the paired heavy-light chain model, the input is a concatenation of heavy and light sequences separated by a special token. Token type IDs are set to 0 for the ‘CLS’ token, 1 for the heavy chain amino acids and 2 for the light chain amino acids to identify two types of chains. Position IDs are set to be the integer position of the amino acid within its respective chain. The Pfam language model was initialized randomly. All other language models were initialized with the pre-trained Pfam model. For all models, the learning rate is set to 10^−5^, batch size is 1024 and the warm-up step is 10,000. One training epoch is defined as one full iteration over all the sequences in the training data. All models were trained until convergence of the cross-entropy loss value (which is evaluated on the validation data after every epoch), or until the maximum number of epochs,10, was reached. All models were trained on NVIDIA Volta V100 GPUs using a distributed compute architecture.

The standard average perplexity score is used to evaluate the language model performance on the hold-out test data. The perplexity measures how well the trained language models are at predicting the masked tokens. Lower values indicate better performance. The average perplexities of the 4 language models on the respective test data are 13.15 for the Pfam model, 1.56 for the heavy-chain model, 1.43 for the light-chain model and 1.16 for the paired model. When evaluated on the OAS light-chain test data, the average perplexities of the 4 language models are 7.47, 16.40, 1.43 and 1.42, respectively. When evaluated on the OAS heavy-chain test data, the average perplexities of the 4 language models are 12.20, 15.30, 1.56 and 1.56, respectively.

### Experimental Binding Measurements for Sequence-to-Affinity Model Training

We separately published the training data to support ease-of-reuse [23]. Briefly, experimental measurements were made, in technical triplicate, by A-Alpha Bio, LLC and are a “predicted affinity” value [27]. This value is derived by comparing the number of binding interactions observed between a pair of proteins to the number of binding interactions of protein-protein pairs with known affinity. When a replicate had no binding measurement reported, we assumed poor binding. For this work, we trained sequence-to-affinity models for the Ab-14-H heavy-chain and Ab-14-L light-chain variants. Each dataset was randomly split into train/validation/test sets with 0.8/0.1/0.1 split.

### Training Sequence-to-Affinity Models via Transfer Learning

We trained separate target-specific sequence-to-affinity models for Ab-14-H variants and Ab-14-L variants. We used model fine-tuning as a way to transfer knowledge learned from pre-trained language models to predicting sequence affinities. We investigated two approaches, which in addition to affinity prediction, provide estimates of prediction uncertainties: an ensemble method and Gaussian Process (GP). Both approaches use learned knowledge from pretrained language models and provide meaningful sequence-to-affinity models from which one can design a diverse antibody library.

The ensemble model consists of 16 different trained regression models that were fine-tuned from the 4 pretrained language models with two different regression loss functions and two different data preprocessing steps (Supplementary Table 5). The two loss functions used were the mean squared error (MSE) and the mean absolute error (MAE) between the predicted affinities and measured affinities. For the data preprocessing step, we considered two options for how we treated missing values, i.e., antibodies with blank binding measurements. Since the experimental assay on the initial antibody library was conducted in triplicate (each antibody sequence has 3 binding measurements), we either drop the assay with missing value or impute it with the median value of all assays of the same candidate chain. Then we take the average of all binding measurements corresponding to the same antibody. To train a regression model, we fine-tuned the pre-trained BERT language model (initially trained on massive sequence data without affinity measurements) by continuing to train it on a smaller set of antibody sequences with experimental binding measurements. The outputs of the ensemble model are the mean and the standard deviation of the outputs of the 16 regression models.

While the ensemble method is known to enhance predictive performance, GP is another powerful technique used for quantifying uncertainties. For the GP model, We used the pretrained heavy-chain language model to train the GP model for the heavy chain sequence-to-affinity model and the pretrained light-chain language model for the light chain sequence-to-affinity GP model. Sequences were represented by first concatenating the learned vector representations of each amino acid from the pretrained language model, and then performing principal component analysis (PCA) to reduce the vector dimension to 1,024. The GP model was trained on these reduced vector representations. The trained GP model outputs a mean and a standard deviation of the binding affinity prediction.

### ML-Extrapolated Fitness Functions

To generate a high affinity scFv library in silico, we used a Bayesian-based acquisition function extrapolated from the sequence-to-affinity model to construct the scFv fitness landscape. In contrast to non-Bayesian settings where the sequence is mapped directly to estimated affinity, the fitness function is defined to be a mapping from the entire scFv sequence to a posterior probability f(x) = p(aff(x)<σ|x) that the estimated binding affinity aff(x) of the sequence x is better than a threshold. The threshold was set to the averaged assayed value of Ab-14 in the training data. Assuming a Gaussian distribution, f(x) can be computed using the mean and standard deviation of the prediction from the trained sequence-to-affinity model. For each scFv chain (Ab-14-H and Ab-14-L), we computed two fitness functions, extrapolated from the ensemble model and GP model, respectively. The proposed fitness function captures the model uncertainty during the optimization and enables us to estimate the performance of our antibody designs prior to experimental testing.

### Optimization Strategies via Sampling

The goal is to sample scFv sequences with the highest extrapolated fitness value f(x). The optimization was performed using 3 different sampling algorithms: a greedy algorithm called hill climb (HC) [28], an evolutionary algorithm called genetic algorithm (GA) [29] and Gibbs sampling [30]. We initialized the HC and GA sampling processes with the 10 strongest binders (seed sequences) from the supervised training data and the Gibbs sampling with the strongest binders from the training data.

For the hill climb algorithm, we initialized the optimization by randomly mutating a seed sequence with an expected number of k=2 mutations. At each step, the algorithm performs a local search around the current sequence and samples the next sequence that has the highest fitness value. The search continues until it can no longer find a sequence that has a better fitness value than the current sequence. We defined the local search space to be the 1,000 mutants of the current sequence, consisting of all the k=1 mutations and random k=2 mutations. The greedy-based hill climb was run 100 times with random restart around a random seed sequence.

The genetic algorithm (GA) is an evolution-based search heuristic, where the fittest individuals are selected to produce offspring of the next generation. We initialized the population with a random seed sequence from the top 10 binders. Parents were chosen from the current population based on the Wright-Fisher model of evolution [31] where members of the current population become parents with a probability exponential to their fitness values, that is, p(x)~exp(f(x)/β). Sequences with high fitness have more chances to pass their genes to the next generation. A single-point crossover was performed on two parent sequences randomly selected from the parent population, and followed by randomly mutating individual child sequences with an expected k=1 mutation. The algorithm was terminated when it no longer produced new sequences (the population converged). The algorithm was run 100 times; each was initialized from a random seed sequence. The parameter *β* was set to be 0.2 for the ensemble-based fitness function and 0.5 for the GP-based fitness function. The selection of parameter value *β* directly affects the diversity of generated sequence designs. Depending on the design needs, one can tune this parameter to adjust the overall library design.

Gibbs sampling is a Markov Chain Monte Carlo (MCMC) algorithm that samples a sequence according to some joint distribution by generating random variates from each of the full conditional distributions. We initialized the algorithm from the top seed sequence (the sequence with the strongest binding affinity in the training data). At each step, we randomly selected a position *i* in the sequence, sampled a mutant 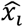 at the selected position with a conditional probability *p*(*x_i_*|*x*_1_ … *x*_*i*−1_, *x*_*i*+1_…*x_L_*) and updated the sequence by replacing the *i^th^* token with the sampled token 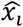. The conditional probability was defined to be exponential to the fitness values, that is, *p*(*x_i_*|*x*_1_ … *x*_*i*−1_, *x*_*i*+1_…*x_L_* ~exp(γ*f(x)). The Gibbs sampling was run once with 30,000 iterations. The value γ was set to be 18 for the Ab-14-H ensemble-based fitness function, and 20 for both the Ab-14-L ensemble- and GP-based fitness function. Multiple γ values were used to sample the Ab-14-H GP-based fitness function. This is due to the limited number of sequences that can be sampled at any specific γ value for the given fitness function. To ensure that enough sequences can be sampled, we used γ=10,3,2 and ran the Gibbs algorithm three times to sample a sufficient number of sequences.

### ML-Optimized ScFv Libraries

For each scFv chain (Ab-14-H variants and Ab-14-L variants), we constructed two fitness functions extrapolated from the ensemble and GP model, respectively. For each fitness function, we performed optimization using three sampling strategies. This resulted in 6 libraries per chain: 3 libraries from optimizing the ensemble-based fitness function (namely, En-HC, En-GA and En-Gibbs), and 3 libraries from optimizing the GP-based fitness function (namely, GP-HC, GP-GA, GP-Gibbs). We then rank-ordered the generated sequences based on their fitness score per library and selected the top 6,000 sequences per library for experimental validation. Supplementary Fig. 2 and Fig. 3 show the distribution of the designed sequences with respect to various mutational distances to demonstrate the library diversity: (1) mutational distance to the candidate scFv Ab-14, (2) mutational distance to the nearest top 10 binders and (3) pairwise mutational distance in a library. The first two distance metrics measure the number of mutations the designed antibodies are from Ab-14 and the best sequences in the training data, respectively. The third distance metric measures the intra-library diversity.

### Evolution Directed Libraries

We built two baseline libraries based on conventional directed evolution strategies: random mutations and the PSSM-based method. The random mutation library was constructed by randomly mutating amino acid tokens from the seed sequences in the training data with a k=2 average number of mutations. Using this method, 2,097 Ab-14-H heavy-chain variants and 477 Ab-14-L light-chain variants were generated for experimental testing.

For the PSSM-based library, we used sequences in the training data with measured affinities that are as good or better than the candidate scFv Ab-14. We fitted the PSSM by counting the occurrence of each amino acid at each position in the CDRs with a small pseudocount. The fitted PSSM is a matrix of probability scores for each amino acid at each position, representing the statistical patterns of the training sequences that are better than Ab-14. We then drew samples to generate designs based on the fitted PSSM. Contrary to the random mutation approach, the PSSM-based approach is not restricted to a pre-defined mutational distance and could generate sequences that are far from the candidate antibody if the computed PSSM allows. The PSSM method resulted in 7,748 Ab-14-H heavy-chain variant designs and 8,257 Ab-14-L light-chain variant designs that were sent for experimental testing. Supplementary Fig. 2 and Fig. 3 show the distribution of the generated sequences with respect to the mutational distances.

### Experimental Validation of the Designed Sequences

The AlphaSeq assay was performed by A-Alpha Bio LLC, with the target protein, scFv libraries and additional positive and negative controls to accommodate the large library sizes. The AlphaSeq assay was conducted from six technical replicates. Some in silico library members are absent from the resulting empirical dataset because they were unsuccessfully mapped in the haploid step and thus had no binding affinity data available. Supplementary Table 6 and Table 7 summarize the number and percentage of sequences present in the experimental data.

For evaluation, we only consider designs that are present in the experimental AlphaSeq data. For sequences that are present in the affinity data and have at least three out of six empirical affinity values, the values are averaged and used as ground-truth measured affinities. Sequences with two or fewer empirical measurements are considered poor binders, and are included in the performance evaluation as un-successful designs. Supplementary Fig. 4 shows the binding distribution of selected libraries and the ML-optimized library outperforms the directed evolution approaches.

### Biophysical property calculation, statistical analysis of libraries, and UMAP embedding

Biophysical properties were calculated based on the sequences of the heavy and light chain variants in each library using BioPython [32]. Specifically, isoelectric points were calculated using *pK* values and methods described by [33]–[35]. Hydrophobicity was calculated using the Kyte & Doolittle index [36]. The hydrophobicity score of each amino acid was averaged over the sequence of each variant to give an overall hydrophobicity score for each sequence. These properties were compared between libraries using the independent 2 sample t-test implemented in scipy [37] with default parameters.

## Acknowledgements

We would like to thank the scientists at A-Alpha Bio for assistance in generation of data; Eric Schwoebel, Joshua Dettman, and Tim Lu for thoughtful discussion of the research program and approach; Jack McGowan and Irene Stapleford for graphic design support; and the many other colleagues at MIT LL who have supported this project.

## Author Contributions

R.C., T.B., and M.W., conceived the project; R.C. and M.W. designed the data collection process; L.S. performed in silico mutagenesis; L.L., R.C., and T.B. designed model architectures; L.L., E.W., J.S., and T.B. performed algorithm development; L.L., E.W., J.S., and L.S. trained models; L.L., E.W., and L.S. evaluated models; L.L., E.W., J.S., T.B., and M.W. performed data analysis; L.L., R.J., R.C., T.B., and M.W. wrote and edited the manuscript.

## Data availability

Raw data for the training data has been deposited to Zenodo at DOI:

## Code availability

Code for training self-supervised language models and supervised sequence-to-affinity models are available on github

