## Supplementary Material for "Machine Learning Optimization of Candidate Antibodies Yields Highly Diverse Sub-nanomolar Affinity Antibody Libraries"

### Supplementary Information

**Supplementary Table 1. Target sequence and candidate antibody sequences (CDRs in bold).** Three candidate antibodies (i.e., Ab-14, Ab-91, Ab-95) were first identified *in vitro* from a human-derived naive phage-display campaign against the target peptide. Two heavy chains (H) and two light chains (L) from the three antibodies were selected for the generation of the initial antibody library. They are Ab-14-H, Ab-91-H, and Ab-14-L, Ab-95-L.

| **Target** | PDVDLGDISGINAS |
|---|---|
| **Candidate Chains** | **Antibody Sequences** |
| Ab-14-H | EVQLVETGGGLVQPGGSLRLSCAAS**GFTLNSYGIS**WVRQAPGKGPEWVSV**IYSDGRRTFYGDSV**KGRFTISRDTSTNTVYLQMNSLRVEDTAVYYCAK**GRAAGTFDS**WGQGTLVTVSS |
| Ab-14-L | DVVMTQSPESLAVSLGERATISC**KSSQSVLYESRNKNSVA**WYQQKAGQPPKLLIY**WASTRES**GVPDRFSGSGSGTDFTLTISSLQAEDAAVYYC**QQYHRLPLS**FGGGTKVEIK |
| Ab-91-H | EVQLVESGGGLVQPGRSLRLSCAAS**GFTFDDYAMH**WVRQAPGKGLEWVS**GISWNSGSIGYADSVK**GRFTISRDNAENSLYLQMNSLRAEDTALYYCAK**VGRGGGYFDY**WGQGTLVTVSS |
| Ab-91-L | QAVLTQPSSLSASPGASVSLTC**TLRSGINVGTYRIY**WYQQKPGSPPQYLLR**YKSDSDKQQGSGV**PSRFSGSKDASANAGILLISGLQSEDEADYYC**MIWHSSAWV**FGGGTKLTVL |
| Ab-95-H | EVQLVESGAEVKKPGASVKVSCKAS**GYTFTSYGIS**WVRQAPGQGLEWMGW**ISAYNGNTNYAQ**KLQGRVTMTTDTSTSTAYMELRSLRSDDTAVYYCAR**VGRGVIDH**WGQGTLVTVSS |
| Ab-95-L | SSELTQDPAVSVALGQTVRITC**EGDSLRYYYAN**WYQQKPGQAPILVIY**GKNNRPS**GIADRFSGSNSGDTSSLIITGAQAEDEADYYC**SSRDSSGFQVF**FGAGTKLTVL |

**Supplementary Table 2. Distribution of mutations within each initial antibody library.** Given the candidate antibody chain, *k*-point mutations, where *k= 1, 2,* and *3*, were designed across all CDRs of each chain. Point mutations were limited to amino acid substitutions.

| Library | k=1 | k=2 | k=3 |
|---|---|---|---|
| Ab-91-H Variants | 684/521 | 4,141/3131 | 25,075/18820 |
| Ab-14-H Variants | 665/594 | 4,089/3671 | 25,146/22188 |
| Ab-14-L Variants | 627/552 | 3,982/3491 | 25,291/22180 |
| Ab-95-L Variants | 551/548 | 3,755/3743 | 25,594/25526 |

**Supplementary Table 3. Train, validation and test splits for protein/antibody language model training.** Values in the parenthesis indicate the number of heavy-light sequence pairs in OAS.

| Datasets | Train | Validation | Test |
|---|---|---|---|
| Pfam | 32,593,668 | 1,715,454 | 44,311 |
| OAS Heavy Chains | 172,524,747 | 47,603,347 | 51,043,837 |
| OAS Light Chains | 70,059,824 | 364,332 | 414,635 |
| OAS Heavy-Light Chains | 242,612,962 (28,391) | 47,972,437 (4758) | 51,459,204 (732) |

**Supplementary Table 4. Regression models used in the ensemble-based fitness model.**

| Name | Base Model | Loss Function | Missing Values |
|---|---|---|---|
| pfam_drop_l1 | Pfam language model | MAE | Drop |
| pfam_drop_mse | Pfam language model | MSE | Drop |
| pfam_median_l1 | Pfam language model | MAE | Impute with median value |
| pfam_median_mse | Pfam language model | MSE | Impute with median value |
| heavy_drop_l1 | Heavy-chain model | MAE | Drop |
| heavy_drop_mse | Heavy-chain model | MSE | Drop |
| heavy_median_l1 | Heavy-chain model | MAE | Impute with median value |
| heavy_median_mse | Heavy-chain model | MSE | Impute with median value |
| light_drop_l1 | Light-chain model | MAE | Drop |
| light_drop_mse | Light-chain model | MSE | Drop |
| light_median_l1 | Light-chain model | MAE | Impute with median value |
| light_median_mse | Light-chain model | MSE | Impute with median value |
| paired_drop_l1 | Paired model | MAE | Drop |
| paired_drop_mse | Paired model | MSE | Drop |
| paired_median_l1 | Paired model | MAE | Impute with median value |
| paired_median_mse | Paired model | MSE | Impute with median value |

**Supplementary Table 5. Percentage incorporation of Ab-14-H designs by library.** Some sequences are absent from the resulting experimental dataset because they were unsuccessfully mapped in the haploid step and thus had no binding affinity data available.

| Library | No. Sequences Designed | No. Sequences Present | Overall % Present |
|---|---|---|---|
| Ensemble-HC | 6000 | 5344 | 89% |
| Ensemble-Gen | 6000 | 5310 | 89% |

| | | | |
|---|---|---|---|
| Ensemble-Gibbs | 6000 | 4879 | 81% |
| GP-HC | 6000 | 5152 | 86% |
| GP-Gen | 6000 | 5313 | 89% |
| GP-Gibbs | 6000 | 5284 | 88% |
| PSSM | 7748 | 6510 | 84% |

**Supplementary Table 6. Percentage incorporation of Ab-14-L designs by library.** Some sequences are absent from the resulting experimental dataset because they were unsuccessfully mapped in the haploid step and thus had no binding affinity data available.

| Library | No. Sequences Designed | No. Sequences Present | Overall % Present |
|---|---|---|---|
| Ensemble-HC | 6000 | 5962 | 99% |
| Ensemble-Gen | 6000 | 5960 | 99% |
| Ensemble-Gibbs | 6000 | 5950 | 99% |
| GP-HC | 6000 | 5965 | 99% |
| GP-Gen | 6000 | 5989 | 100% |
| GP-Gibbs | 6000 | 5987 | 100% |
| PSSM | 8257 | 8188 | 99% |

**Supplementary Table 7. The best antibody heavy chain sequences by library**

| Libraries | Best Ab-14-H Variant |
|---|---|
| Random Mutations | [redacted] |
| PSSM | [redacted] |
| GP-HC | [redacted] |
| GP-Gen | [redacted] |
| GP-Gibbs | [redacted] |
| En-HC | [redacted] |
| En-Gen | [redacted] |
| En-Gibbs | [redacted] |

| | |
|---|---|
| | [redacted] |

**Supplementary Table 8. The best antibody light chain sequences by library**

| Libraries | Best Ab-14-L Variant |
|---|---|
| Random Mutations | [redacted] |
| PSSM | [redacted] |
| GP-HC | [redacted] |
| GP-Gen | [redacted] |
| GP-Gibbs | [redacted] |
| En-HC | [redacted] |
| En-Gen | [redacted] |
| En-Gibbs | [redacted] |

**Supplementary Table 9. Ranking of Libraries Using Estimated Percent of Success**

| Ensemble Models | | | | | GP Models | | | | |
|---|---|---|---|---|---|---|---|---|---|
| Method | Ab-14-H Variant Libraries | | Ab-14-L Variant Libraries | | Method | Ab-14-H Variant Libraries | | Ab-14-L Variant Libraries | |
| | Rank (Predicted) | Rank (Actual) | Rank (Predicted) | Rank (Actual) | | Rank (Predicted) | Rank (Actual) | Rank (Predicted) | Rank (Actual) |
| PSSM | 4 | 3 | 4 | 4 | PSSM | 4 | 4 | 4 | 4 |
| En-HC | 2 | 2 | 1 | 1 | GP-HC | 3 | 3 | 2 | 3 |
| En-GA | 1 | 1 | 2 | 3 | GP-GA | 1 | 1 | 1 | 1 |
| En-Gibbs | 3 | 4 | 3 | 2 | GP-Gibbs | 2 | 2 | 3 | 2 |
| | Rank Correlation: 0.8 | | Rank Correlation: 0.8 | | | Rank Correlation: 1 | | Rank Correlation: 0.8 | |

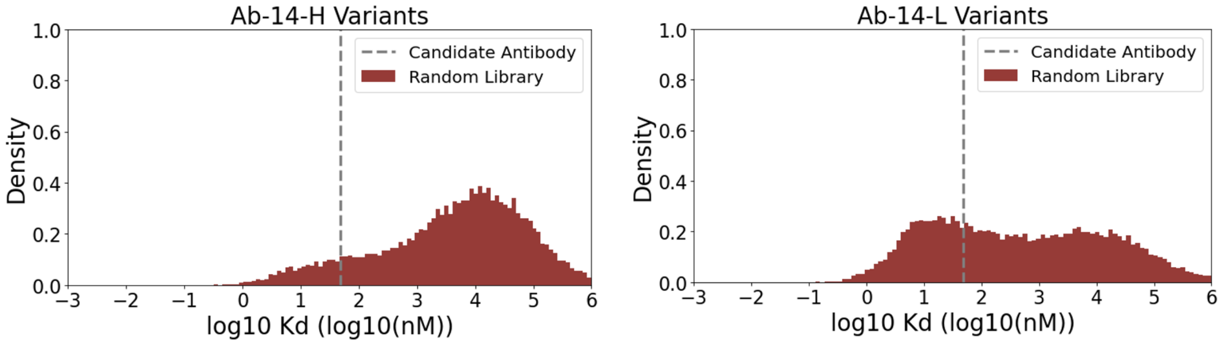

**Supplementary Fig. 1 Library distribution for Ab-14-H and Ab-14-L variants.** Average affinity is used for each sequence with at least 2 (out of 3) measured binding values.

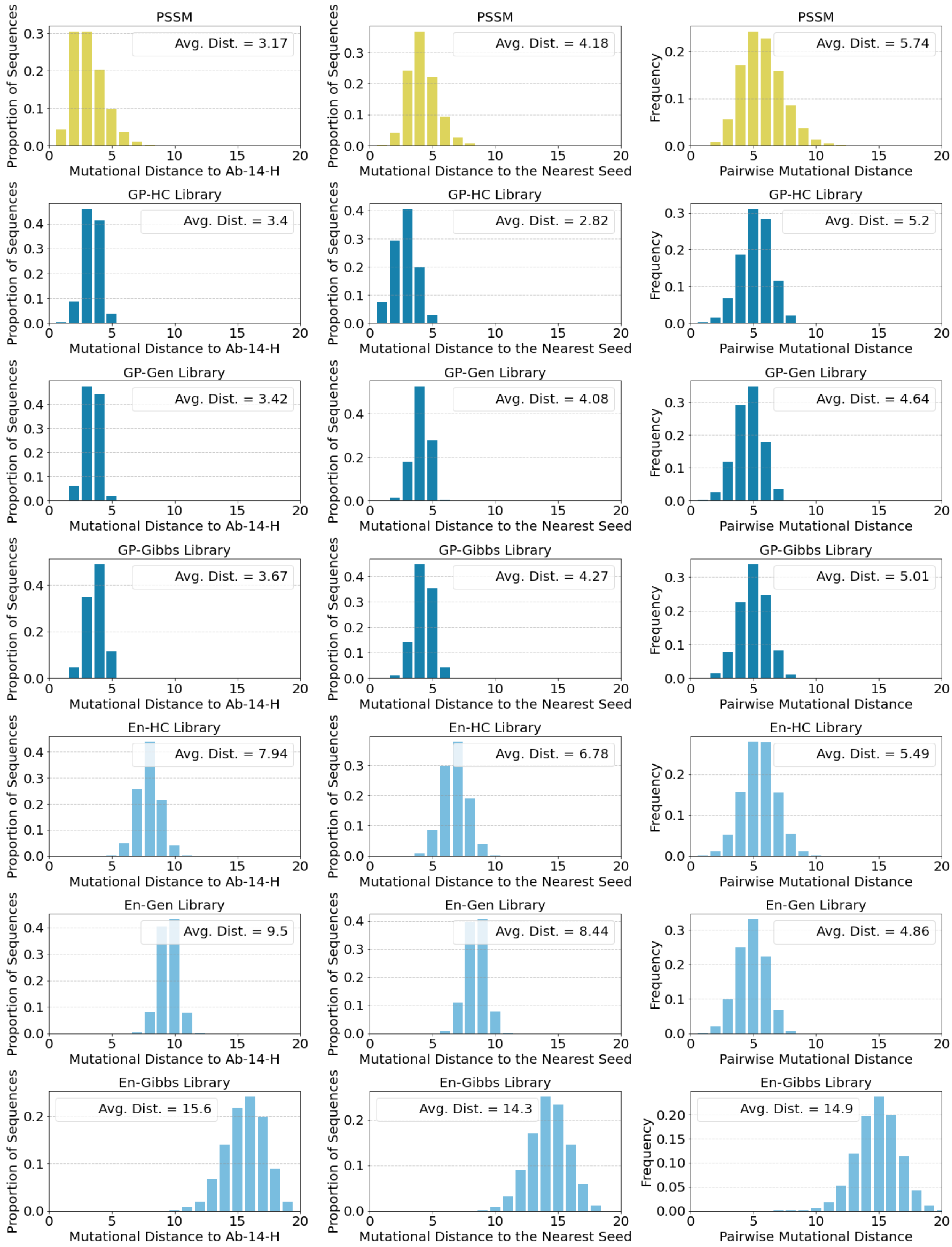

**Supplementary Fig. 2 Distribution of diversity metrics by library for Ab-14-H variants**

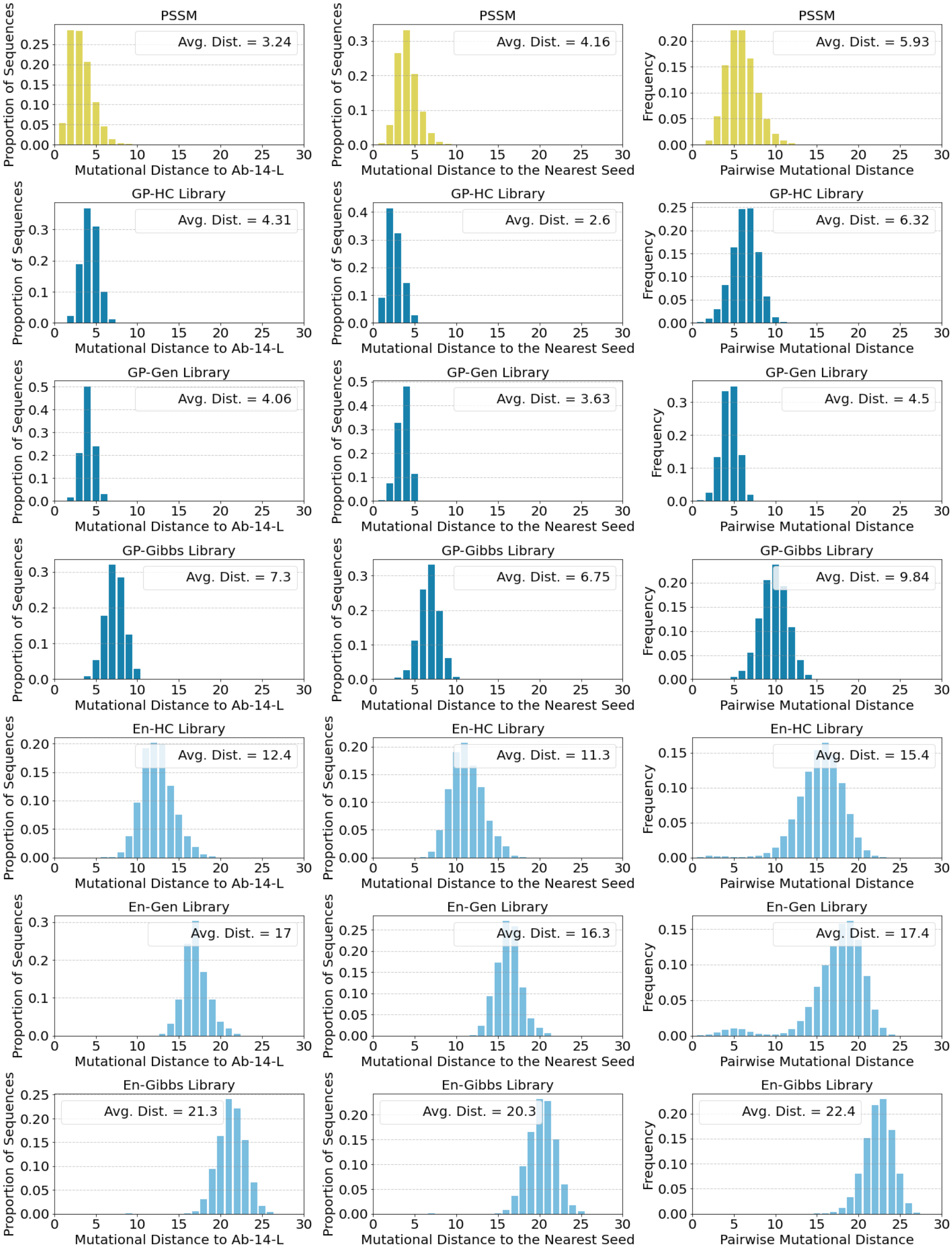

**Supplementary Fig. 3 Distribution of diversity metrics by library for Ab-14-L variants**

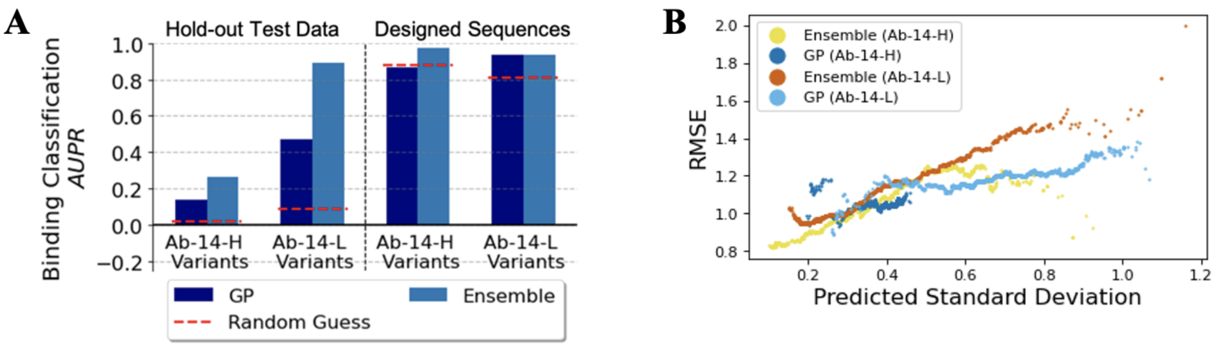

**Supplementary Fig. 4 Sequence-to-affinity model evaluation.** (A) AUPR of classification task (strong binders vs weak binders); AUPR is tailored for the detection of rare events as there is very few number of strong binders in the hold-out test data from the training set. (B) Evaluation of the uncertainty estimate of sequence-to-affinity models and the root mean squared error (RMSE) of the prediction.

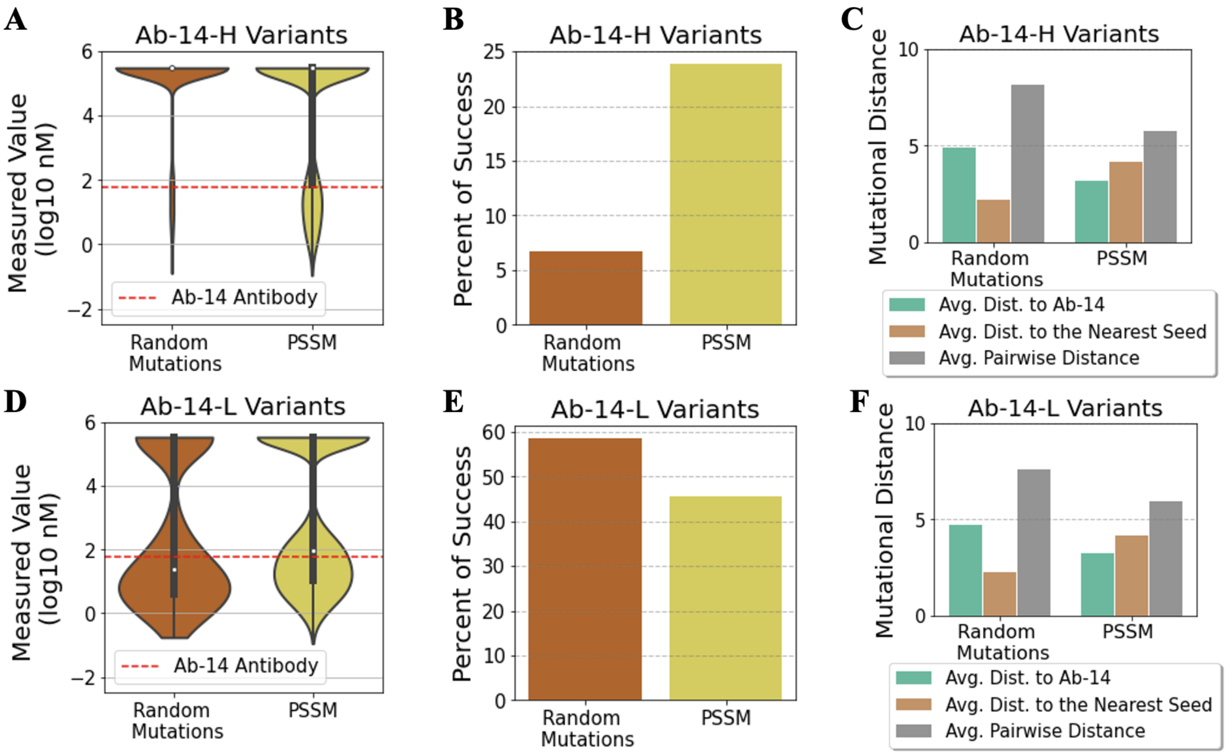

**Supplementary Fig. 5 Random mutation library vs PSSM library.** The binding quantification assay was conducted from six technical replicates. A replicate with an empty measured affinity value indicates that it was beyond the limit of detection and was deemed a poor binders. For sequences that have at least 3 (out of 6) empirical binding affinities, we use the averaged values as ground-truth measured affinities. The rest of the sequences (with less than 3 empirical measurements) are automatically considered as un-successful designs. (A) Measured affinity distribution of Ab-14-H heavy chain designs (excluding sequences with no ground-truth affinity value). Affinities of unsuccessful sequences are set to be 5.48 (the largest assay value of all Ab-14-H variants). (B) Percent of sequences that have stronger binding affinity than the candidate antibody for all the Ab-14-H variant libraries. (C) Diversity comparison for all the Ab-14-H variant libraries. (D) Measured affinity distribution of Ab-14-L light chain designs. Affinities of unsuccessful sequences are set to be 5.53 (the largest assay value of all Ab-14-L variants). (E) Percent of success for all the Ab-14-L variant libraries. (F) Diversity comparison for all the Ab-14-L variant libraries.

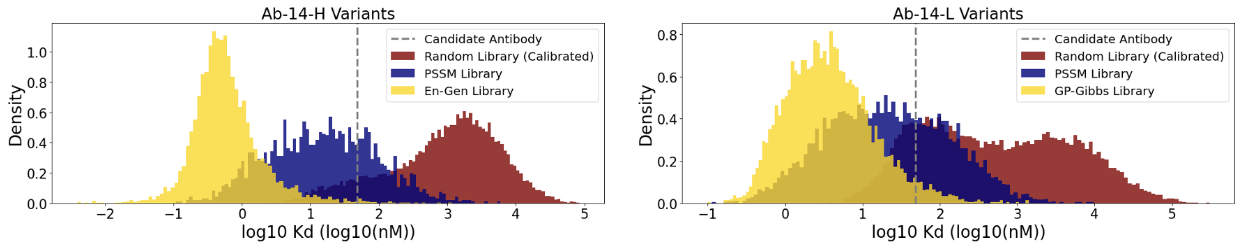

**Supplementary Fig. 6 Distribution of designed antibodies on selected libraries.** The affinity values for the random library are calibrated to match the distribution of the affinities over the common sequences.

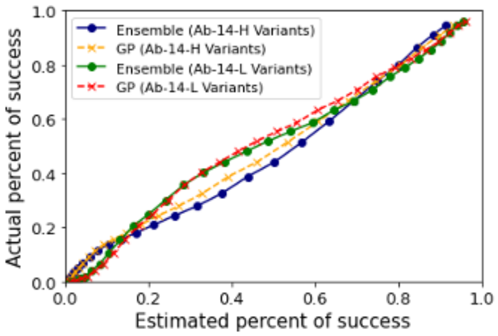

**Supplementary Fig. 7 Evaluation of proposed in silico metric for library performance prediction.** The estimated percent of success aligned well with the true performance on the hold-out test data.

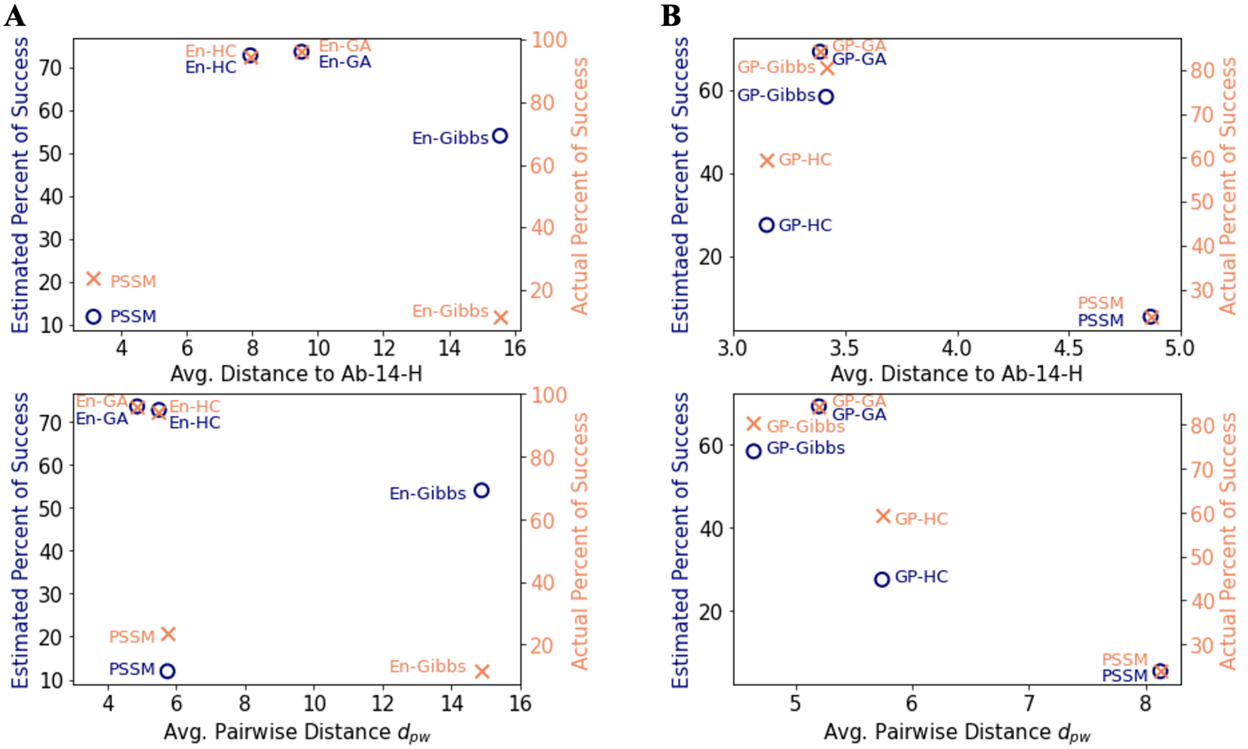

**Supplementary Fig. 8 The estimated percent of success metric enables exploration of the tradeoffs between performance and diversity and informs library selection (Ab-14-H variant designs).** The relative performance of the estimated percent of success matches well to the empirically measured percent of success. (A) ensemble-based libraries for Ab-14-H variants. (B) GP-based libraries for Ab-14-H variants.

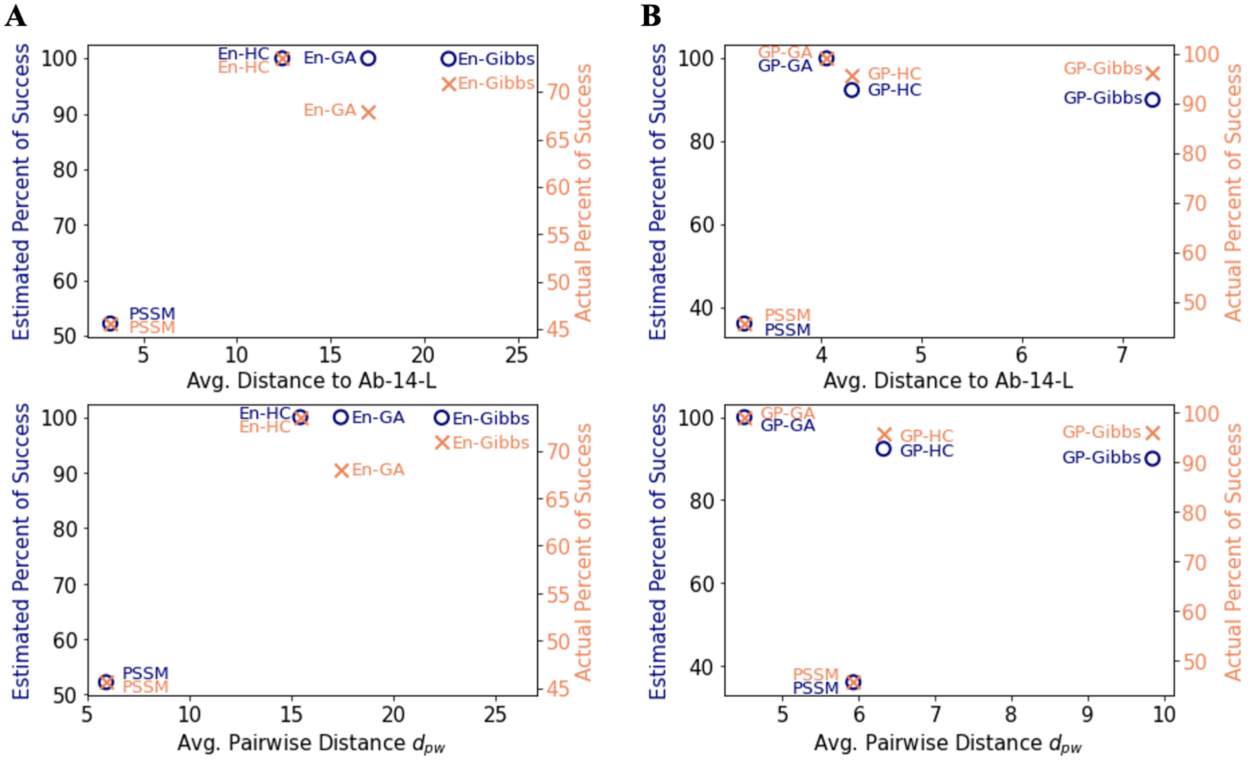

**Supplementary Fig. 9 The estimated percent of success metric enables exploration of the tradeoffs**

**between performance and diversity and informs library selection (Ab-14-L variant designs).** The relative performance of the estimated percent of success matches well to the empirically measured percent of success. (A) ensemble-based libraries for Ab-14-L variants. (B) GP-based libraries for Ab-14-L variants.

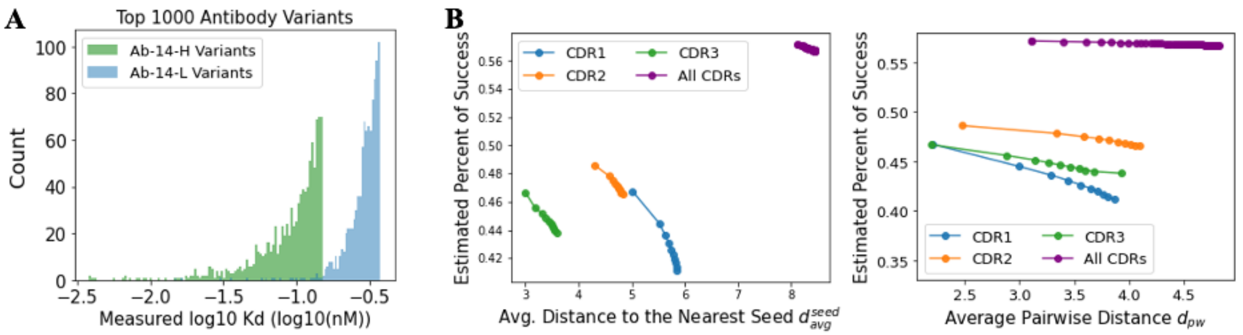

**Supplementary Fig. 10 Evaluation of designing various antibody CDR regions.** (A) Histogram of top 1000 heavy chain and light chain designs; heavy chain designs led to significantly stronger binders. (B) Library comparison on designing the entire heavy chain CDRs and individual CDRs as the number of sequences in each library increases. Designing the entire heavy chain CDRs produces antibodies with higher estimated percent of success and diversity.

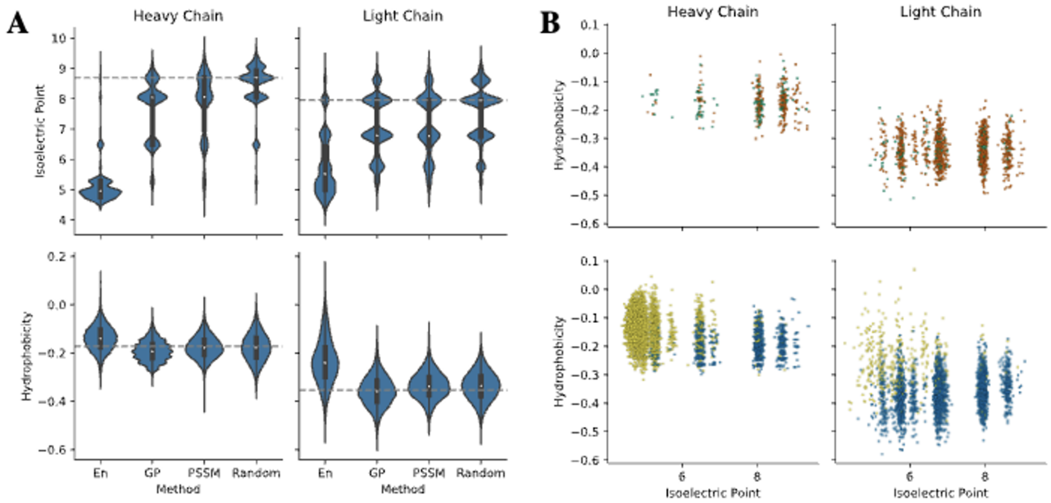

**Supplementary Fig 11. Biophysical and sequence properties of heavy and light chain libraries.** (A) Violin plots of isoelectric points and hydrophobicities of heavy and light chain sequences in the random library and the PSSM, GP, and ensemble designed libraries. (B) Scatter plots showing the joint distribution of isoelectric points and hydrophobicities for sequences with strong binding affinity (measured binding affinity <= 1 nM). Orange: random, green: PSSM, blue: GP, and yellow: ensemble-designed sequences. Our designed strong binders cover a wide range of these biophysical properties.
